# Connecting conceptual and spatial search via a model of generalization

**DOI:** 10.1101/258665

**Authors:** Charley M. Wu, Eric Schulz, Mona M. Garvert, Björn Meder, Nicolas W. Schuck

## Abstract

The idea of a “cognitive map” was originally developed to explain planning and generalization in spatial domains through a representation of inferred relationships between experiences. Recently, new research has suggested similar principles may also govern the representation of more abstract, conceptual knowledge in the brain. We test whether the search for rewards in conceptual spaces follows similar computational principles as in spatial environments. Using a within-subject design, participants searched for both spatially and conceptually correlated rewards in multi-armed bandit tasks. We use a Gaussian Process model combining generalization with an optimistic sampling strategy to capture human search decisions and judgments in both domains, and to simulate human-level performance when specified with participant parameter estimates. In line with the notion of a domain-general generalization mechanism, parameter estimates correlate across spatial and conceptual search, yet some differences also emerged, with participants generalizing less and exploiting more in the conceptual domain.

## Introduction

The ability to search for rewards comes in many shapes. We can wander through a foreign city in search of new and delicious foods, or search through an online store to find a laptop with the features that we like. We can even skim over parts of a paper to find sections more interesting than the introduction. While these tasks differ in a number of ways, all of them require the exploration of possibilities and the use of generalization to predict outcomes of unexplored options. Here, we ask if generalization and search in different domains can employ common computational mechanisms.

Breaking from the classical stimulus-response school of reinforcement learning, Tolman (1948) argued that both rats and humans extract a cognitive representation from experience, described as a “cognitive map” of the environment. Rather than merely representing stimulus-response associations, cognitive maps also encode inferred relationships between experiences or options, such as the distances between locations in space, thereby facilitating planning ahead and generalization. While cognitive maps were first identified as representations of physical spaces, Tolman also hypothesized that similar principles may underlie the organization of knowledge more broadly (Tolman, 1948).

The idea of a cognitive map has been widely adopted in research on brain signals underlying spatial navigation (O’Keefe & Nadel, 1978). In a similar vein, studies on reinforcement learning have emphasized that relations between states may also be encoded as a cognitive map (Schuck, Cai, Wilson, & Niv, 2016; Sutton & Barto, 1998). Most recently, a number of studies suggest that the same neural representations may underlie the organization of spatial and non-spatial relational information in the brain (Constantinescu, O’Reilly, & Behrens, 2016; Garvert, Dolan, & Behrens, 2017; Kaplan, Schuck, & Doeller, 2017). This is consistent with behavioral evidence for generalized cognitive search processes (Hills, Todd, & Goldstone, 2008). Whereas the capacity to *generalize* from past experiences to unobserved states and actions has been studied for decades (Shepard, 1987), the link between our ability to generalize in a wide range of tasks and the encoding of experiences in a cognitive map-like format has not yet been explored.

To assess this link, we investigate how people search for rewards in both spatially and conceptually correlated multi-armed bandit tasks (Stojic, Analytis, & Speekenbrink, 2015; Wu, Schulz, Speekenbrink, Nelson, & Meder, 2017). In one task, the spatial correlation of rewards can be used to guide generalization based on the spatial similarity between options. In the other task, participants can use feature-based similarity to navigate conceptual space in the search for rewards. In both tasks, search takes place in state spaces larger than the available search horizon and with noisy rewards, thereby inducing an exploration-exploitation dilemma. Generalization is constrained by the level of environmental correlation, which we vary between participants.

Our results show that participants are able to learn and generalize in both tasks, with performance correlated across spatial and conceptual domains. Surprisingly, performing the spatial task first boosted performance in the conceptual task, but not vice versa. We apply computational modeling to understand how spatial and conceptual environments may lead to differences in how people generalize about unobserved rewards and how they approach the exploration-exploitation dilemma. Using a computational model based on *Gaussian Processes* (GP) combined with *Upper Confidence Bound* (UCB) sampling, we predict participant choices and judgments in both search domains, and achieve human-level performance using participant parameter estimates. We find that participant parameter estimates describing the level of generalization and exploration were correlated across spatial and conceptual domains, but that participants tended to generalize less and exploit more in the conceptual domain.

## Experiment

Participants searched for rewards in two successive multi-armed bandit tasks with correlated rewards (Fig. 1). In one task, rewards were spatially correlated *(Spatial task)*, meaning options with similar spatial locations yielded similar rewards. In the other task, rewards were conceptually correlated *(Conceptual task)*, such that options with similar features (i.e., the number of leaves ∈ [1, 5] and berries ∈ [1, 5]) yielded similar rewards. Figure 1c,d shows examples of fully revealed environments representing the same underlying reward function, but mapped to either spatial or conceptual features. In both tasks, the search space was represented by a 5 × 5 two-dimensional grid, where each of the 25 options represented a different arm of the bandit, which could be clicked to obtain (noisy) rewards. Each tile of the grid contained one of 25 unique conceptual stimuli, which were randomly shuffled between rounds and always visible. Thus, we presented information about both spatial and conceptual features in both search tasks, but only one of them was relevant for generalization and predicting rewards. At the beginning of each round only a single randomly chosen option was revealed (i.e., displayed the numerical reward and corresponding color aid), whereby subjects had a limited horizon of 10 actions in each round (40% of the total search space; similar to Wu et al., 2017), thereby inducing an exploration-exploitation trade-off.

**Figure 1:**
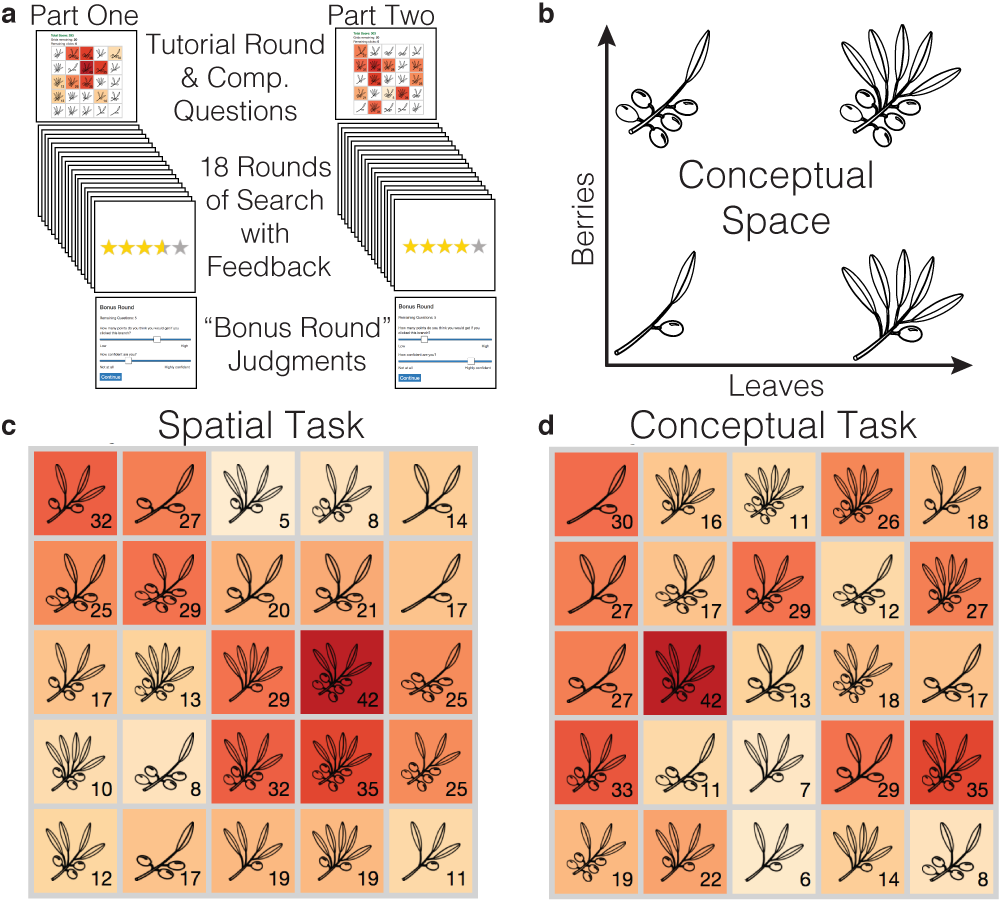
Experiment. **a**) Searching for spatially or conceptually correlated rewards in a two part multi-armed bandit experiment, where task order was counter-balanced across subjects. **b**) Illustration of conceptual space, with four edge cases shown. **c, d**) Two fully revealed environments (representing the same underlying reward function), represented as a 5 × 5 grid, where participants could click on the tiles to obtain rewards, revealing a numeric payoff value and a corresponding color aid (larger rewards are darker; unexplored tiles are initially white). **c:** Spatial search task, where rewards are spatially correlated, with nearby tiles yielding similar rewards. **d:** Conceptual search task, where options with high feature similarity (i.e., number of leaves and number of berries) had similar rewards.

### Methods

**Participants and Design**. 72 participants were recruited through Amazon Mechanical Turk (requiring 95% approval rate and 100 previously approved HITs) for a two part experiment, where only those who completed part one were invited for part two. In total, 64 participants completed both parts of the experiment and were included in the analyses (26 Female; mean age=34, SD=11). Participants were paid $1.25 for each part of the experiment, with those completing both parts being paid an additional performance-contingent bonus of up to $3.00. Participants earned $4.94 ± 0.29 and spent 26 ± 13 minutes completing both parts. There was an average gap of 5.6 ± 4.7 hours between the two parts of the experiment.

Task order varied between subjects, with participants completing the Spatial and Conceptual task in counterbalanced order in separate sessions. We also varied between subjects the extent of reward correlations in the search space by randomly assigning participants to one of two different classes of environments *(Smooth* vs. *Rough)*, with smooth environments corresponding to stronger correlations, and the same environment class used for both tasks.

**Materials and Procedure**. Each search task comprised 20 rounds (i.e., grids), with a different reward function sampled without replacement from the set of assigned environments. The reward function specified how rewards mapped onto either the spatial or conceptual features. Between rounds, the locations of each stick stimuli were randomly shuffled. In each round, participants had a limited search horizon of 10 available actions (i.e., clicks), which could be used to either explore unrevealed options or to exploit known options. Participants were instructed to accumulate as many points as possible, which were later converted into monetary payoffs.

For both tasks, the first round was an interactive tutorial and the last round was a “bonus round” (Fig. 1a). In the tutorial round, participants were shown instructions for the given task alongside an interactive grid, which functioned identically to subsequent rounds. Participants were told that options with either similar spatial features (Spatial task) or similar conceptual features (Conceptual task) would yield similar rewards. Three comprehension questions (different for spatial and conceptual tasks) were used to ensure full understanding of the task (specifically whether spatial or conceptual features predicted reward) before participants were allowed to continue. In the bonus round, participants made explicit judgments about the expected rewards and their estimated uncertainty of five unrevealed tiles in the middle of the round (i.e., after five clicks), in order to tap into beliefs supported by generalization. All behavioral and computational modeling analyses exclude the first and last rounds, except for the analysis of the bonus round judgments.

**Spatial and Conceptual Search Tasks**. At the beginning of each round, one random tile was revealed (i.e., showing numerical payoff and color aid) and participants could click any of the 25 tiles in the grid until the search horizon of ten clicks was exhausted, including re-clicking previously revealed tiles. Clicking an unrevealed tile displayed the numerical value of the reward along with a corresponding color aid, where darker colors indicated higher point values. Previously revealed tiles could also be re-clicked, although there were variations in the observed value due to noise. Each observation included normally distributed noise, 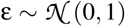, where the rewards for each round were scaled to a uniformly sampled maximum value in the range of 35 to 45, so that the value of the global optima could not be easily guessed. For repeat clicks, the most recent observation was displayed numerically, while hovering over the tile would display the entire history of observations. The color of the tile corresponded to the mean of all previous observations.

Participants were awarded up to five stars based on their performance at the end of each round (e.g., 4.4 out of 5), based on the ratio of their average reward to the global maximum. The performance bonus (up to $3.00) was calculated based on the average number of stars earned in each round, excluding the tutorial round.

**Judgments**. In both tasks the last round was a “bonus round”, which solicited judgments about the expected reward and estimated uncertainty of five unrevealed options. Participants were informed that the goal of the task remained the same (maximize cumulative rewards), but that after five clicks, they would be asked to provide judgments about five randomly selected options, which had not yet been explored (sampled uniformly from unexplored options). Judgments about expected rewards were elicited using a slider, which changed the displayed value and color of the selected tile from 0 to 50 (in increments of 1). Judgments about uncertainty were elicited using a slider from 0 to 10 (in increments of 1), with the endpoints labeled ‘Not at all’ and ‘Highly confident’. After providing the five judgments, participants were asked to choose one of the five selected options to reveal, and subsequently completed the round like all others.

**Environments**. All environments were sampled from a GP prior parameterized with a *radial basis function* (RBF) kernel (see below for details), where the length-scale parameter (λ) determines the rate at which the correlations of rewards decay over (spatial or conceptual) distance. Higher λ-values correspond to stronger correlations. We generated 40 samples of each type of environments, using λ*_Smooth_* = 2 and λ*_Rough_* = 1, which were sampled without replacement and used as the underlying reward function in each task.

### Modeling Generalization and Exploration

We use a combination of a *learning model* with a *decision strategy* to make predictions about each individual participant’s search decision. The learning model forms beliefs about the expectations of rewards *μ*(x) and the associated uncertainty *σ*(**x**) for each option **x**, which are then used by the decision strategy to make probabilistic predictions about search decisions. We apply leave-one-round-out cross validation to estimate the free parameters of our models and use out-of-sample model predictions to compare models for predicting human search behavior. Additionally, we compare predictions of the learning models to judgments made by participants about the expected reward and estimated uncertainty of five unrevealed options.

### Learning models

**Function Learning**. We use Gaussian Process (GP) regression (Rasmussen & Williams, 2006; Schulz, Speekenbrink, & Krause, 2017) as a *Function Learning* model for inducing an underlying value function mapping the features of the search task onto rewards, as a method for generalization. A 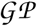 defines a distribution *P*(*f*) over possible functions *f* (**x**) that map inputs **x** to output *y*. In our case, either the spatial features (i.e., *x*- and *y*-coordinates on the grid) or conceptual features (i.e., number of leaves and berries) of each option serve as inputs **x** to predict reward y. Crucially, learning a value function by using either spatial or conceptual similarity allows for predictive generalization of unobserved options (see Wu, Schulz, Speekenbrink, Nelson, & Meder, 2018).

A 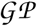 is completely defined by a mean function *m*(**x**) and a kernel function, *k*(**x**, **x**′):

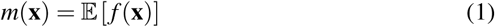

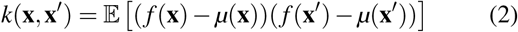

We fix the prior mean to the median value of payoffs, *m*(**x**) = 25, while the kernel function *k*(**x**, **x**′) encodes prior assumptions (or inductive biases) about the underlying function. Here, we use the *radial basis function* (RBF) kernel:

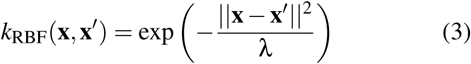

The RBF kernel models similarity by assuming correlations between two options **x** and **x**′ decay as an exponential function of their (spatial or conceptual) distance. The length-scale parameter λ determines how far correlations extend, with larger values of λ assuming stronger correlations over longer distances, whereas λ → 0^+^ assumes complete independence of options. We use recovered parameter estimates of λ to learn about the extent to which participants generalize about unobserved rewards.

**Option Learning**. The *Option Learning* model uses a Bayesian Mean Tracker (BMT) and is an associative learning model (Speekenbrink & Konstantinidis, 2015). In contrast to the GP Function Learning model, the Option Learning model learns the rewards of each option independently by computing independent posterior distributions for the mean *μ_j_* for each option *j*:

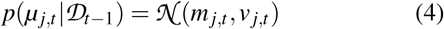

The rewards of each option *j* are learned independently, with the posterior mean *m_j, t_* and variance *v_j, t_* only updated when selected at trial *t*, based on the observed reward *y_t_*:

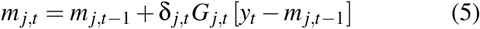

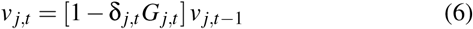

where δ*_j,t_* = 1 if option *j* was chosen on trial *t*, and 0 otherwise. Additionally, the learning factor *G_j,t_* is defined as:

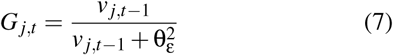

where 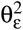 is the error variance, which is estimated as a free parameter. Intuitively, the estimated mean of the chosen option *m_j,t_* is updated based on the prediction error *y_t_ − m_j,t_*_−1_, multiplied by the learning factor *G_j,t_*. At the same time, the estimated variance *v_j,t_* is reduced by a factor of 1 − *G_j,t_*, which is in the range [0, 1]. The error variance 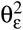 can be interpreted as an inverse sensitivity, where smaller values result in more substantial updates to the mean *m_j,t_*, and larger reductions of uncertainty *V_j,t_*. We set the prior mean to the median value of payoffs *m_j,_*_0_ *=* 25 and the prior variance *v_j,_*_0_ = 250.

### Decision Strategy

Both Function Learning and Option learning models generate normally distributed predictions about the expected reward *μ*(**x**) and estimated uncertainty **σ**(**x**) for each option. These estimates are used by the decision strategy for evaluating the quality *q*(**x**) of each option and making a prediction about where to sample next. We use *Upper Confidence Bound sampling* (UCB) to compute a weighted sum of the expected reward *μ*(**x**) and the estimated uncertainty **σ(x**):

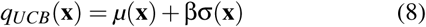

where the exploration factor β determines how the reduction of uncertainty trades off against exploiting high expected rewards; a strategy that has been found to predict search behavior in a variety of contexts (Wu et al., 2018; Schulz, Konstantinidis, & Speekenbrink, 2017).

We then use a softmax function to convert the value of an option *q*(**x**) into a choice probability:

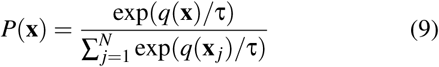

where τ is the temperature parameter. Whereas β encodes exploration directed towards uncertain options, τ encodes undirected (noisy) exploration as a distinct (Wilson, Geana, White, Ludvig, & Cohen, 2014) and separately recoverable (Wu et al., 2018) phenomenon. As τ → 0 the highest-value arm is chosen with a probability of 1 (i.e., argmax); when τ → ∞, predictions converge to random choice.

## Results

### Behavioral Results

Performance was highly correlated between the two tasks (Pearson’s *r* = .74, *p <* .001; Fig. 2a). with rewards being slightly lower in the conceptual task (*t*(63) = −3.7, *p <* .001, *d* = −0.34), although Figure 2b shows how this is largely due to the influence of task order. There were no performance differences between the two tasks when the spatial task was performed first (*t*(34) = 0.6, *p* = .55, *d* = 0.07). However, in the reverse order (conceptual first), mean rewards in the conceptual task were worse than in the spatial task (*t*(28) = −5.6, *p <* .001, *d* = −0.68). Thus, searching first for spatially correlated rewards improved performance in the conceptual domain (*t*(62) = 2.6, *p* = .01, *d* = 0.66), but not vice versa.

**Figure 2:**
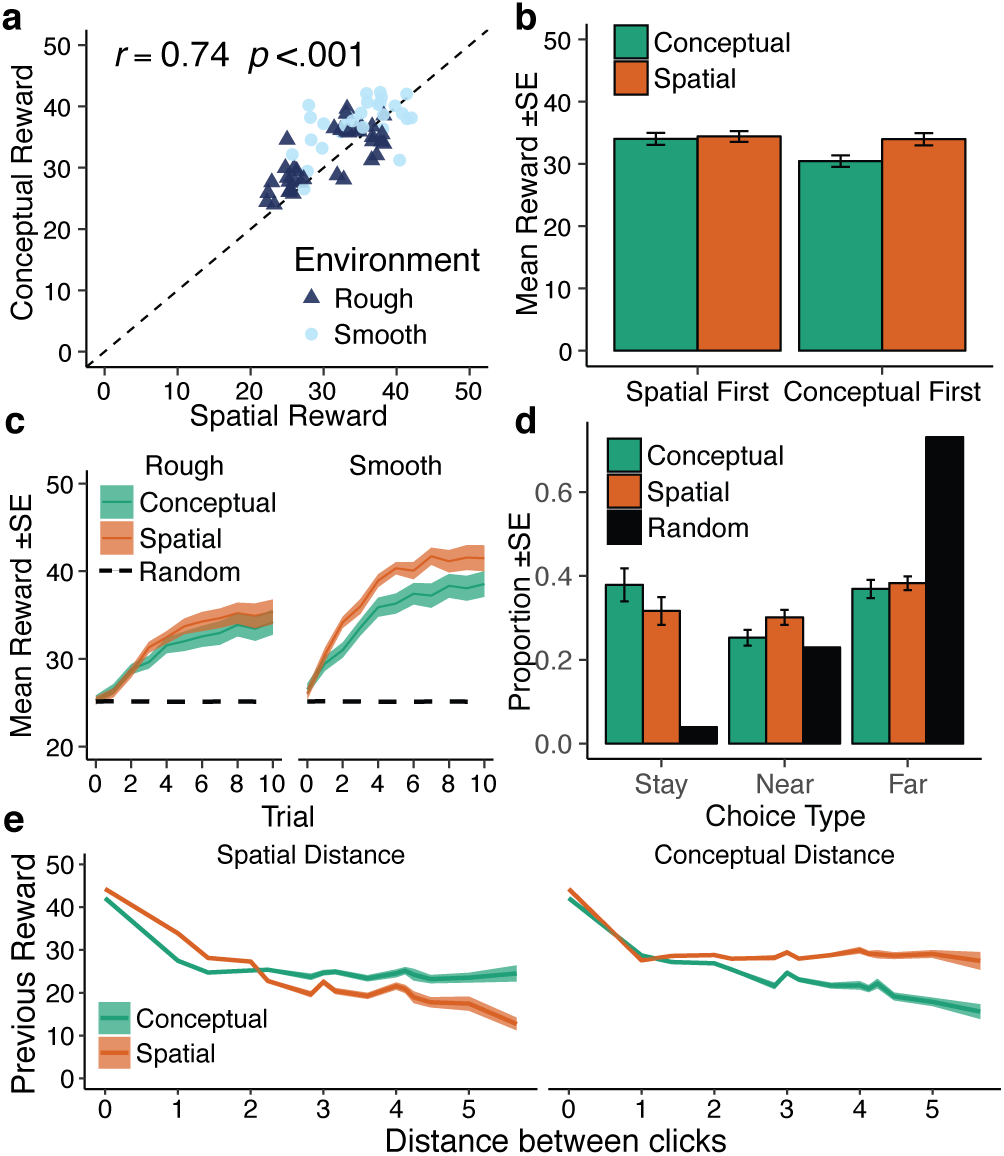
Behavioral results. **a**) Performance in the spatial and the conceptual tasks were correlated, where each point is a single participant, with the dashed line indicating *y* = *x*. **b**) Comparing performance across the counter-balanced task order, we found performing the spatial task first boosted performance on the conceptual task. **c**) Learning curves over trials with comparison to a random baseline (10k replications), showing the mean (line) and standard error (ribbon) aggregated over rounds. d) Proportion of choices based on the distance between clicks with a comparison to a random baseline (10k replications). “Stay” indicates repeat clicks, “Near” indicates a neighboring tile (measured in either spatial or conceptual distance), and “Far” indicates all other possibilities. **e**) Relationship between the value of the previous reward and the Euclidean distance to the next selected option (left: spatial distance; right: conceptual distance), showing mean (line) and standard error (ribbon).

The learning curves in Figure 2c show that participants systematically found higher rewards over subsequent trials (*r* = .51, *p* < .001), performed better in smooth than in rough environments (*t*(62) = 4.4, *p* < .001, *d* = 1.1), and that the performance gap between spatial and conceptual performance was larger in smooth environments. Looking only at participants assigned to smooth environments, performance was better in the spatial task than in the conceptual task (*t*(27) = 3.2, *p* = .003, *d* = 0.59), consistent with the larger gap between learning curves in Figure 2c. We did not find systematic improvements over rounds (*r* = .02, *p* = .42).

**Search Distance** Figure 2d shows the proportion of different types of search decisions, where *stay* corresponds to a repeat click (**x***_t_* = **x***_t_*_−1_), *near* corresponds to searching one of the neighboring options in either spatial or in conceptual distance (i.e., ±1 leaf and/or ±1 berry in feature space), and *far* corresponds to all other possible distances. Participants in the conceptual task tended to make more repeat clicks (stay; *t*(63) = 2.4, *p* = .02, *d* = 0.21), whereas participants in the spatial task were more likely to search neighboring options (near; *t*(63) = 3.4, *p* = .001, *d* = 0.33). In both contexts, participants clearly behaved differently than the random baseline model. Looking at the relationship between the value of a reward and the distance searched on the subsequent trial (Fig. 2e), we see that participants responded appropriately, with a stronger influence of reward value on spatial distance in the spatial task, and a stronger influence of reward on conceptual distance in the conceptual task. This suggests that participants used information from the relevant dimension (spatial or conceptual) to make their decisions.

### Modeling Results

We first compared models based on their ability to predict participants’ behavior using leave-one-round-out cross validation (Fig. 3a), where the *Conceptual GP* and the *Spatial GP* utilize either conceptual or spatial features, respectively. In the spatial task, the Spatial GP performed better than both the Conceptual GP (i.e., using leaf and berry features; *t*(63) = 8.5, *p <* .001, *d* = 0.61) and the BMT (*t*(63) = 4.6, *p* < .001, *d* = 0.36) replicating previous findings reported in Wu et al. (2018). Surprisingly, all models performed equally well in the conceptual task (*F*(2, 189) = 0.27, *p* = .76).

**Figure 3:**
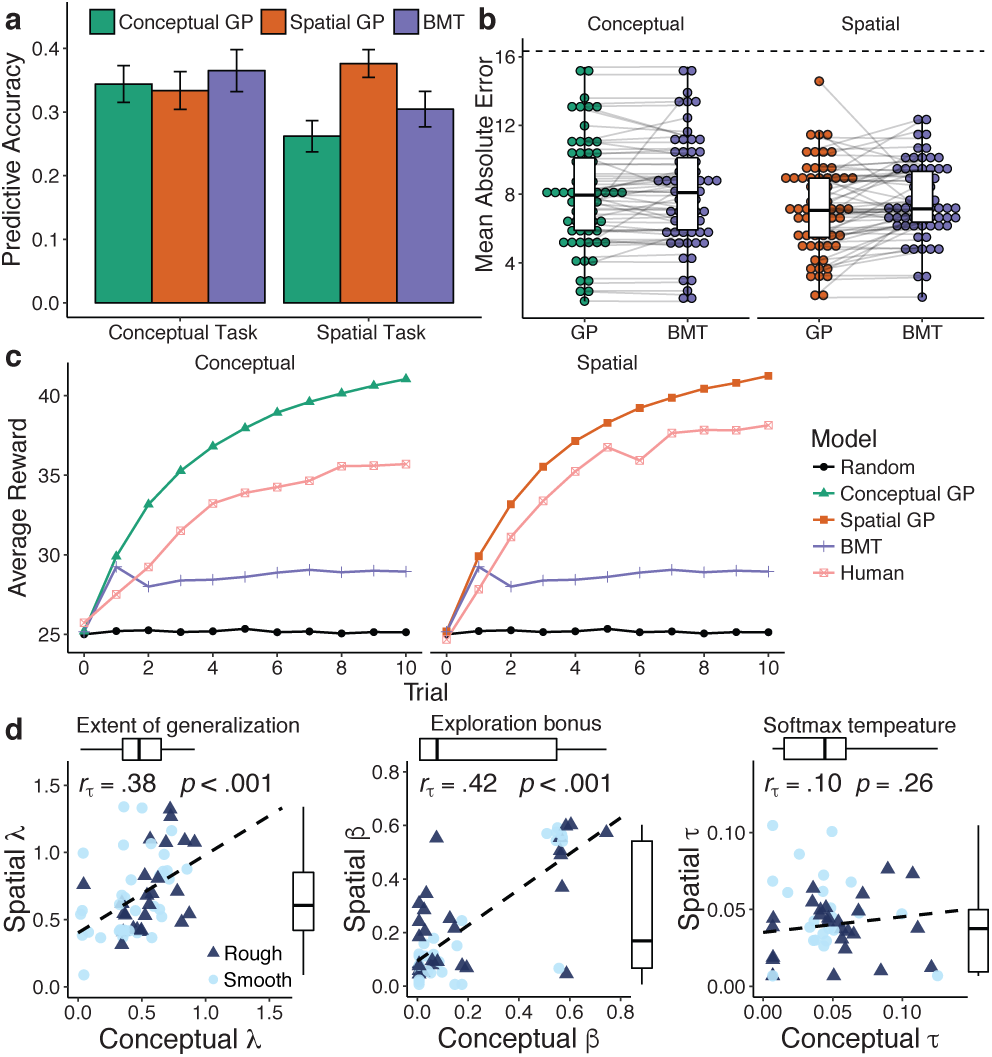
Modeling results. **a**) Predictive accuracy of models on out-of-sample predictions, where 0 corresponds to random chance and 1 is a theoretically perfect model. Bars show mean across participants with error bars indicating standard error. **b**) Mean Absolute Error (MAE) on the judgment task, where lines connect each individual participant (dots) and show the difference in prediction error between the two models. Boxplots show the median and 1.5 IQR. Dashed line shows theoretical MAE of random guesses. **c**) Simulated learning curves (10k replications; aggregated over environments) using models specified with parameters sampled from participant estimates. We include comparison to a random model (black line) and human performance (pink). **d**) Median GP parameters from the spatial (y-axis) and the conceptual tasks (x-axis), where each point is a single participant and the dotted line shows a linear regression. Outliers are excluded from the plot but not from the rank correlations and boxplots (showing median and 1.5 IQR range).

The correspondence between participants’ judgments from the bonus round and model predictions are shown in Figure 3b, where the corresponding GP had lower error than the BMT in the spatial task (*t*(63) = −2.2, *p* = .03, *d* = −0.2), but there was no difference in the conceptual task (*t*(63) = −0.9, *p* = .35, *d* = −0.05). GP predictions were also correlated with participant judgments about expected reward in both the spatial task (*r* = .38, *p* < .001) and the conceptual task (*r* = .21, *p* < .001), whereas the BMT invariably predicted both a mean and variance of 25, making correlations undefined. GP predictions about perceived uncertainty were weakly rank-correlated with participants’ judgments in the conceptual task (Kendall’s rank correlation; *r_τ_* = .12, *p* = .003), but not in the spatial task (*r_τ_* = −.01, *p* = .75).

Importantly, we simulated model performance on the task over 10, 000 replications, where model parameters were sampled from the cross-validated participant estimates. Looking at the simulated learning curves (Fig. 3c), the GP parameter estimates produced human-like performance for both tasks, whereas the BMT performed only marginally better than a random sampling model. Thus, even though the BMT model produces decent predictions, it is not able to produce human-like learning curves. Thus, the simulated learning behavior falsifies the BMT model as a plausible account of human behavior (see Palminteri, Wyart, & Koechlin, 2017).

**Parameter Estimates** Looking at the parameter estimates of the GP for the two different tasks, we find that λ (extent of generalization) and β (exploration bonus) were strongly rank-correlated within participants (Fig. 3d), whereas the softmax temperature *τ* was not correlated and also did not differ between tasks (Z = 1.1, *p* = .13, *r* = .14). Interestingly, although λ-values were correlated across the two tasks, they were significantly smaller in the conceptual task than in the spatial task (Wilcoxon signed rank test; *Z* = −5.1, *p* < .001, *r* = .64), meaning participants generalized over smaller conceptual distances than over spatial distances. We also found that larger lambdas were correlated with higher performance across both tasks (*r*_τ_ = .45, *p* < .001). Estimates for the exploration factor β did not differ between tasks (Z = 1.1, *p* = .86, *r* = .14). Thus, participants who generalized more or displayed more directed exploration in one task, also did so in the other, connecting both spatial and conceptual search.

## General Discussion and Conclusion

Humans search for rewards across a multitude of different domains, using effective generalization and clever exploration to great success. Historic psychological findings explained adaptive generalization in spatial domains by evoking the concept of a cognitive map, whereas more recent neuroscientific evidence suggests cognitive maps can be found in both spatial and conceptual domains. We investigated whether the search for spatially or conceptually correlated rewards can be connected via common principles of generalization. Our results showed that participants performed well in both domains, with highly correlated performance across the two tasks. Using a Gaussian Process regression framework as a model of generalization and postulating Upper Confidence Bound sampling as an optimistic approach to the exploration-exploitation dilemma, we made progress towards understanding the computational mechanisms of generalization and search across spatial and conceptual domains. Our model produced good out-of-sample predictions in both tasks, made predictions of unobserved rewards that correlated with participants’ judgments, and produced human-like learning curves based on meaningful parameter estimates that showed levels of generalization and directed exploration that correlated across the two domains.

Nevertheless, the GP was not able to predict participant choices better than a non-generalizing option learning model in the conceptual domain, even though the behavioral data indicates successful generalization (see Fig. 2c,e). This could be explained by two different—not mutually exclusive— reasons. One explanation is that the conceptual task was simply more difficult, which led participants to generalize less and sample more locally. Another explanation could be that conceptual stimuli induce different priors over features than in the spatial domain. For example, participants may have strongly linear priors for conceptual features (e.g., more berries will lead to higher rewards) or that they have assumptions about the importance of different features (e.g., berries are more important for rewards than leaves). To overcome these problems of prior assumptions, we could directly assess participant priors over different stimuli and specify our models using the resulting empirical priors. To overcome the problem of differentially perceived feature importance, we could assess the performance of kernels with direct relevance determination (Gershman & Daw, 2017), which similar to attentional weights (see Niv et al., 2015), could also be used to predict participant choices.

We explored whether the same model of generalization can be used to explain how people search for rewards in either spatial or conceptual domains. Our results showed that some aspects of human behavior can indeed be explained by shared computational principles, such as the ability to generalize about unobserved outcomes and a tendency to explore uncertain outcomes. Nonetheless, some clear differences emerged, with participants generalizing less and exploiting more in the conceptual domain, and with transfer only occurring unidirectionally from the spatial to the conceptual task (see also Hills et al., 2008). However, the intra-subject consistency of model parameters across the two domains offers an exciting opportunity in the future to use neural imaging to study if there is indeed a common neural basis for how people generalize and explore both spatial and conceptual spaces. We believe that further study into the general principles underlying human generalization and exploration will continue to provide important insights into adaptive behavior in complex and uncertain environments.

## Acknowledgments

CMW is supported by the International Max Planck Research School on Adapting Behavior in a Fundamentally Uncertain World; ES is supported by the Harvard Data Science Initiative; BM is supported by grant ME 3717/2–2 as part of the DFG priority program “New Frameworks of Rationality” (SPP 1516).

